# Ketogenesis is dispensable for the metabolic adaptations to caloric restriction

**DOI:** 10.1101/2025.05.29.656153

**Authors:** Chung-Yang Yeh, Alexis L. Borgelt, Brynn J. Vogt, Alyssa A. Clark, Ted T. Wong, Isaac Grunow, Michelle M. Sonsalla, Reji Babygirija, Yang Liu, Michaela E. Trautman, Mariah F. Calubag, Bailey A. Knopf, Fan Xiao, Dudley W. Lamming

**Affiliations:** Department of Medicine, University of Wisconsin-Madison, Madison, WI 53705, USA; William S. Middleton Memorial Veterans Hospital, Madison, WI 53705, USA; Comparative Biomedical Sciences Graduate Training Program, University of Wisconsin-Madison, Madison, WI 53706, USA; Graduate Program in Cellular and Molecular Biology, University of Wisconsin-Madison, Madison, WI 53706, USA; Nutrition and Metabolism Graduate Program, University of Wisconsin-Madison, Madison, WI 53706, USA; Endocrinology and Reproductive Physiology Graduate Program, University of Wisconsin-Madison, Madison, WI 53706, USA; University of Wisconsin Comprehensive Diabetes Center, University of Wisconsin-Madison, Madison, WI, 53705, USA; University of Wisconsin Carbone Comprehensive Cancer Center, University of Wisconsin-Madison, Madison, WI 53705, USA; Wisconsin Nathan Shock Center of Excellence in the Basic Biology of Aging, Madison, WI 53705, USA

**Keywords:** caloric restriction, dietary restriction, metabolism, ketogenesis, ketones, BHB, metabolic health

## Abstract

Caloric restriction (CR) extends the health and lifespan of diverse species. When fed once daily, CR-treated mice rapidly consume their food and endure a prolonged fast between meals. As fasting is associated with a rise in circulating ketone bodies, we investigated the role of ketogenesis in CR using mice with whole-body ablation of *Hmgcs2*, the rate-limiting enzyme producing the main ketone body β-hydroxybutyrate (βHB). Here, we report that *Hmgcs2* is largely dispensable for many metabolic benefits of CR, including CR-driven changes in adiposity, glycemic control, liver autophagy, and energy balance. Although we observed sex-specific effects of *Hmgcs2* on insulin sensitivity, fuel selection, and adipocyte gene expression, the overall physiological response to CR remained robust in mice lacking *Hmgcs2*. To gain insight into why the deletion of *Hmgcs2* does not disrupt CR, we measured fasting βHB levels as mice initiated a CR diet. Surprisingly, as mice adapt to CR, they no longer engage high levels of ketogenesis during the daily fast. Our work suggests that the metabolic benefits of long-term CR are not mediated by ketogenesis.

## Introduction

Caloric restriction (CR) extends healthspan and longevity in a variety of species ranging from yeast, worms, and flies to mice and non-human primates, with benefits for healthy aging seen as well in short-term human studies (Green, Lamming, & Fontana, 2022). CR-treated mice are typically fed a single daily meal containing 20-40% fewer total calories than the energy intake of mice fed a matched *ad libitum* (AL) diet. This style of CR leads to the mice binge eating their daily food allotment in 2-3 hours, self-imposing a prolonged fast between meals (V. A. Acosta-Rodriguez, de Groot, Rijo-Ferreira, Green, & Takahashi, 2017; Bruss, Khambatta, Ruby, Aggarwal, & Hellerstein, 2010; Duregon, Pomatto-Watson, Bernier, Price, & de Cabo, 2021). This daily fasting period contributes to the benefits of CR in both wild-type mice (Pak et al., 2021) and Alzheimer’s disease mice (Babygirija et al., 2025).

Fasting elicits a plethora of physiological adaptations (Mihaylova et al., 2023). One key metabolic response, as blood glucose levels are depleted, is to initiate ketogenesis and produce ketone bodies from fatty acids (de Cabo & Mattson, 2019). Of the three ketone bodies – acetoacetate, acetone, and 3-β-hydroxybutyrate (βHB) – βHB is the most abundant and well-studied. βHB acts both as an energy currency and as a signaling molecule, for instance, through direct post-translational modifications and the modulation of histone deacetylases (Newman & Verdin, 2017). Fasting, ketogenic diets, and dietary supplementation with exogenous ketones can all significantly elevate circulating βHB levels and can elicit benefits in Alzheimer’s disease animal models (Di Lucente et al., 2024; Madhavan et al., 2025; Ye et al., 2024). A critical unanswered question is whether fasting-induced ketogenesis contributes to the beneficial effects of CR (Mihaylova et al., 2023).

In naturally-aged mice, there is an age-dependent decline in ketone body metabolism (Eap et al., 2022), highlighting a potential role of ketone bodies in the aging process. Ketogenesis is mediated by mitochondrial 3-hydroxy-3-methylglutaryl-CoA synthase 2 (HMGCS2), the rate-limiting enzyme in the formation of ketone bodies. HMGCS2 has been shown to drive fatty acid β-oxidation, and promotes expression of the lifespan-modulating energy balance hormone FGF21 (Vila-Brau, De Sousa-Coelho, Mayordomo, Haro, & Marrero, 2011; Zhang et al., 2012). Together, these data support the idea that ketone body metabolism is intricately connected with aging and health-boosting diets that engage ketogenesis.

To determine how ketogenesis contributes to the metabolic response to CR, we generated a mouse model lacking *Hmgcs2*. We placed both male and female *Hmgcs2^-/-^*(KO) mice on either an *ad libitum* (AL) diet or a 30% reduced CR diet (CR), with CR-fed animals receiving food once per day at the beginning of the light cycle. We extensively characterized the metabolic phenotypes of these animals, finding that KO mice broadly received the same benefits from CR as wild-type (WT) mice, with few genotype-diet interactions. To understand why the loss of *Hmgcs2* led to unexpectedly limited consequences to the CR response, we measured the circulating βHB level over 24 hrs. Surprisingly, the WT mice treated with prolonged CR do not produce high levels of βHB as the matching AL mice when fasting. In a separate cohort of mice, we demonstrated that this downregulation in ketogenesis occurs during the first few weeks of CR initiation. Together, our results suggest that ketogenesis is dispensable for the metabolic benefits of CR and that ketone bodies are not elevated during long-term CR due to adaptations to recurrent daily fasting.

## Results

### *Hmgcs2* deletion does not alter the effects of CR on weight and body composition

To investigate the role of ketogenesis in caloric restriction (CR), we generated mice lacking whole body *Hmgcs2* expression in the C57BL/6J genetic background. At 9 weeks of age, male and female Hmgcs2^+/+^ (WT) and Hmgcs2^-/-^ (KO) mice were singly housed and either maintained on an *ad libitum* diet (AL) or “stepped down” to a 30% reduced CR diet, fed once per day within 1 hr from the start of the light cycle (**Fig. 1A**). The CR-fed mice were pair-fed according to the AL group, with meal size adjusted weekly. Food consumption and body weight (BW) were tracked weekly, and metabolic testing began after 8 weeks on diet. A schematic of this workflow is presented in **Supplemental Fig. 1A**. Following the completion of the experiments, we confirmed that the HMGCS2 protein was undetectable in KO mice across multiple tissues, including the liver, small intestine (jejunum and ileum), large intestine (colon), and kidney (**Figs. 1B-1E**). Notably, in WT mice of both sexes, CR strongly induced HMGCS2 expression in the kidney but not in other tissues (**Fig. 1E**).

**Figure 1.**
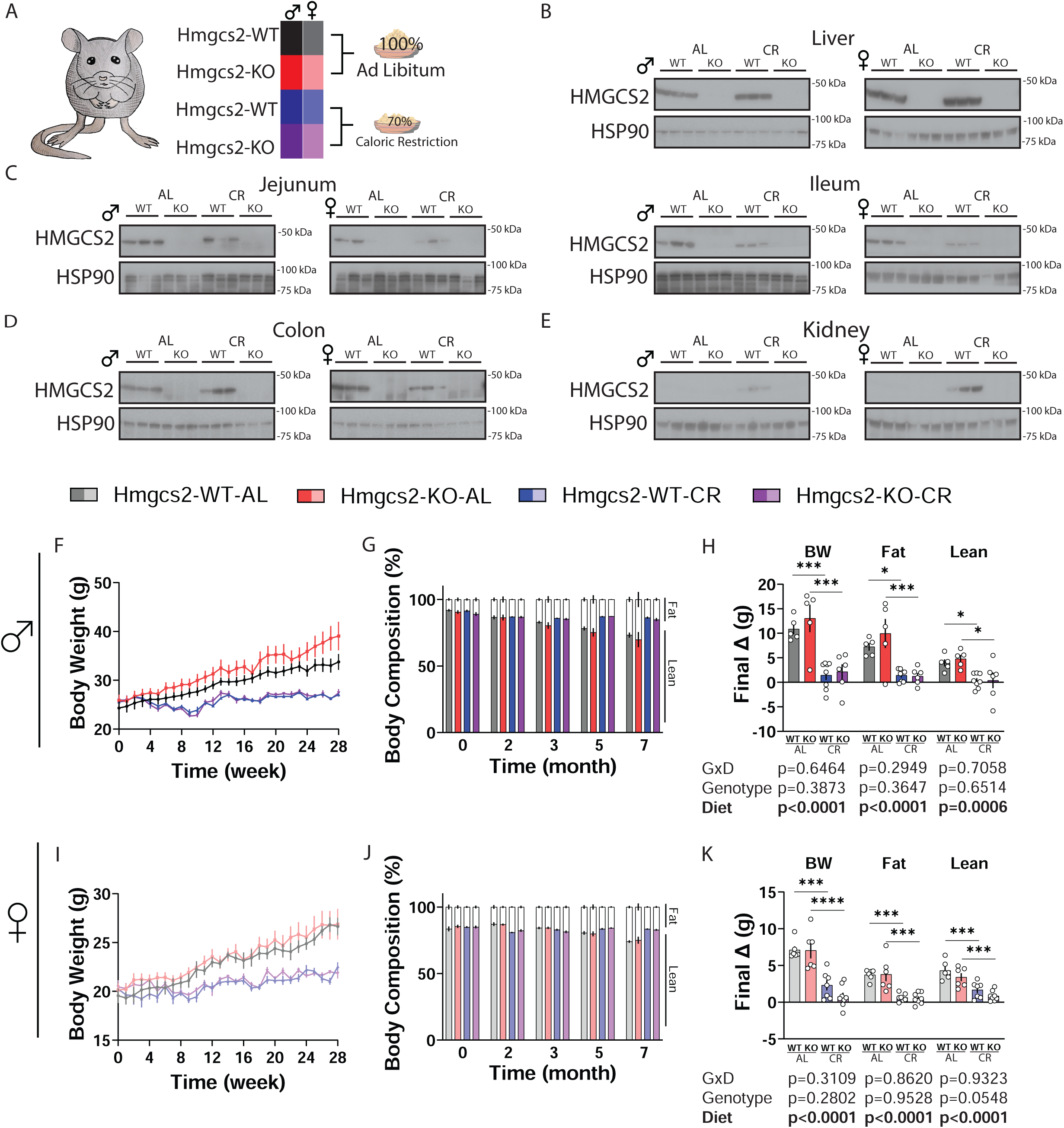
The ablation of ketogenesis does not alter the effect of caloric restriction on body weight and body composition. **(A)** Main experimental setup of the caloric restriction study. **(B-E)** Western blot confirmation of the ablation of Hmgcs2 in the KO animals of both AL and CR groups in the liver (B), the small intestine jejunum and ileum (C), the large intestine colon (D), and the kidney (E). **(F-H)** Male mice body weight (F), body composition % (G), and final changes in body composition (H) over 7 months. For (F) and (G), at the beginning of the experiment, WT-AL, KO-AL, WT-CR, KO-CR, n=7,6,8,7 respectively; for (H) n=5,5,8,6. **(I-K)** Female mice body weight (I), body composition % (J), and final changes in body composition (K) over 7 months. For (I) and (J), at the beginning of the experiment, WT-AL, KO-AL, WT-CR, KO-CR, n=6,7,7,9 respectively; for (K) n=6,6,7,8. (H & K) *p<0.05, ***p<0.001, ****p<0.0001, Sidak’s test post 2-way ANOVA. Data presented as mean ± SEM.

As anticipated, CR suppressed weight gain over a 7-month period in both WT and KO male mice (**Fig. 1F**). Throughout the experiment, the AL group did not exhibit notable changes in food consumption (**Supplemental Fig. 1B**). Knockout of *Hmgcs2* did not affect the ability of CR to suppress the gain of fat (**Supplemental Fig. 1C**) and lean mass (**Supplemental Fig. 1D**), and we observed the corresponding changes in body composition (**Fig. 1G; Supplemental Fig. 1E**). Nearly identical observations were made in the female mice (**Figs. 1I-1J; Supplemental Figs. 1F-1I**). Analysis of final changes in BW, fat, and lean mass revealed only robust diet effects and no significant effects of genotype (**Figs. 1H & 1K**).

### *Hmgcs2* deletion improves insulin sensitivity in female mice

To determine the effects of *Hmgcs2* deletion on glucose metabolism, we performed glucose, insulin, and pyruvate tolerance tests. As our previous work has shown that time since feeding dictates insulin sensitivity in CR (Pak et al., 2024), tests were conducted after either 21 hrs or 7 hrs of fasting, which corresponds to the CR feeding time of approximately 6 am, or 7 hrs after CR feeding at approximately 4 pm, respectively.

We observed an overall positive effect of CR on glucose metabolism. CR improved glucose tolerance in both male and female mice regardless of the length of fasting; however, there was no effect of genotype or a genotype x diet interaction in either sex (**Figs. 2A & 2E**; **Supplemental Figs. 2A & 3A**). In agreement with our previous observations (Pak et al., 2024), CR improved the response to insulin after 21 hrs of fasting in both sexes (**Figs. 2B & 2F**). With the shorter 7 hrs fasting period, this effect on insulin response was reversed, with insulin resistance observed in the CR-fed females (**Supplemental Figs. 2B & 3B**). Interestingly, the deletion of *Hmgcs2* had an overall beneficial effect in females regardless of the length of the fast, significantly increasing insulin sensitivity in female AL mice when insulin sensitivity was assessed after 21 hrs of fasting (**Figs. 2F; Supplemental Fig. 3B**).

**Figure 2.**
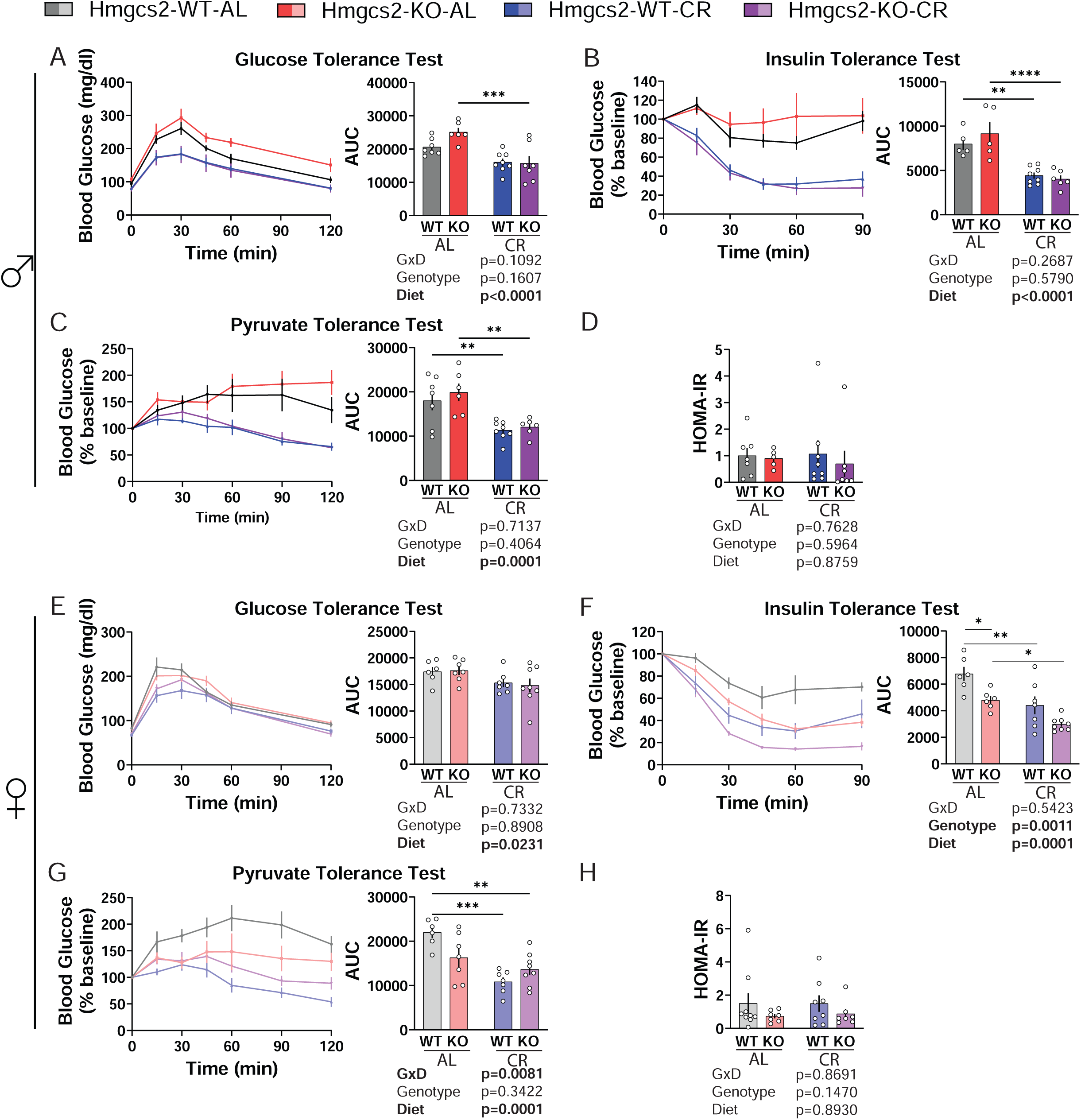
The ablation of ketogenesis selectively improves insulin sensitivity in females. **(A-D)** After 21 hrs fasting in male mice, the results of glucose tolerance test (A; 1 g/kg; I. P.), insulin tolerance test (B; 0.5 U/kg; I. P.), pyruvate tolerance test (C; 2 g/kg; I. P.), and HOMA-IR measurements (D). For WT-AL, KO-AL, WT-CR, and KO-CR in each experiment, GTT n=7,6,8,7; ITT n=7,6,8,6; PTT n=7,6,8,6; HOMA-IR n=7,5,8,7. **(E-H)** After 21 hrs fasting in female mice, the results of glucose tolerance test (E; 1 g/kg; I. P.), insulin tolerance test (F; 0.5 U/kg; I. P.), pyruvate tolerance test (G; 2 g/kg; I. P.), and HOMA-IR measurements (H). For WT-AL, KO-AL, WT-CR, and KO-CR in each experiment, GTT n=6,7,7,8; ITT n=6,7,6,8; PTT n=6,7,7,8; HOMA-IR n=9,7,8,7. (A-H) *p<0.05, **p<0.01, ***p<0.001, ****p<0.0001, Sidak’s test post 2-way ANOVA. Data presented as mean ± SEM.

A pyruvate tolerance test examines the ability of the liver to suppress gluconeogenesis. We found that CR improved pyruvate tolerance in all groups (**Figs. 2C & 2G; Supplemental Figs. 2C & 3C**). In females, although there was no overall effect of genotype, we observed a significant genotype-diet interaction, suggesting that *Hmgcs2* contributes to CR-mediated effects on pyruvate tolerance (**Fig. 2G**).

In a glucose-stimulated insulin secretion test after 21 hrs fasting, there was a trend in the suppression of insulin secretion in the males by CR (p=0.0643) (**Supplemental Figs. 2D and 3D**). There was no overall effect of diet, genotypes, or their interaction on insulin sensitivity as assessed by HOMA-IR in either sex (**Figs. 2D & 2H**). However, there was a trend (p=0.1470) towards improved insulin sensitivity in the KO females (**Fig. 2H**). Overall, while we identified some interesting sex-specific genotype effects and genotype-diet interactions, we did not find the ablation of *Hmgcs2* to significantly impede the main metabolic characteristics of CR.

**Figure 3.**
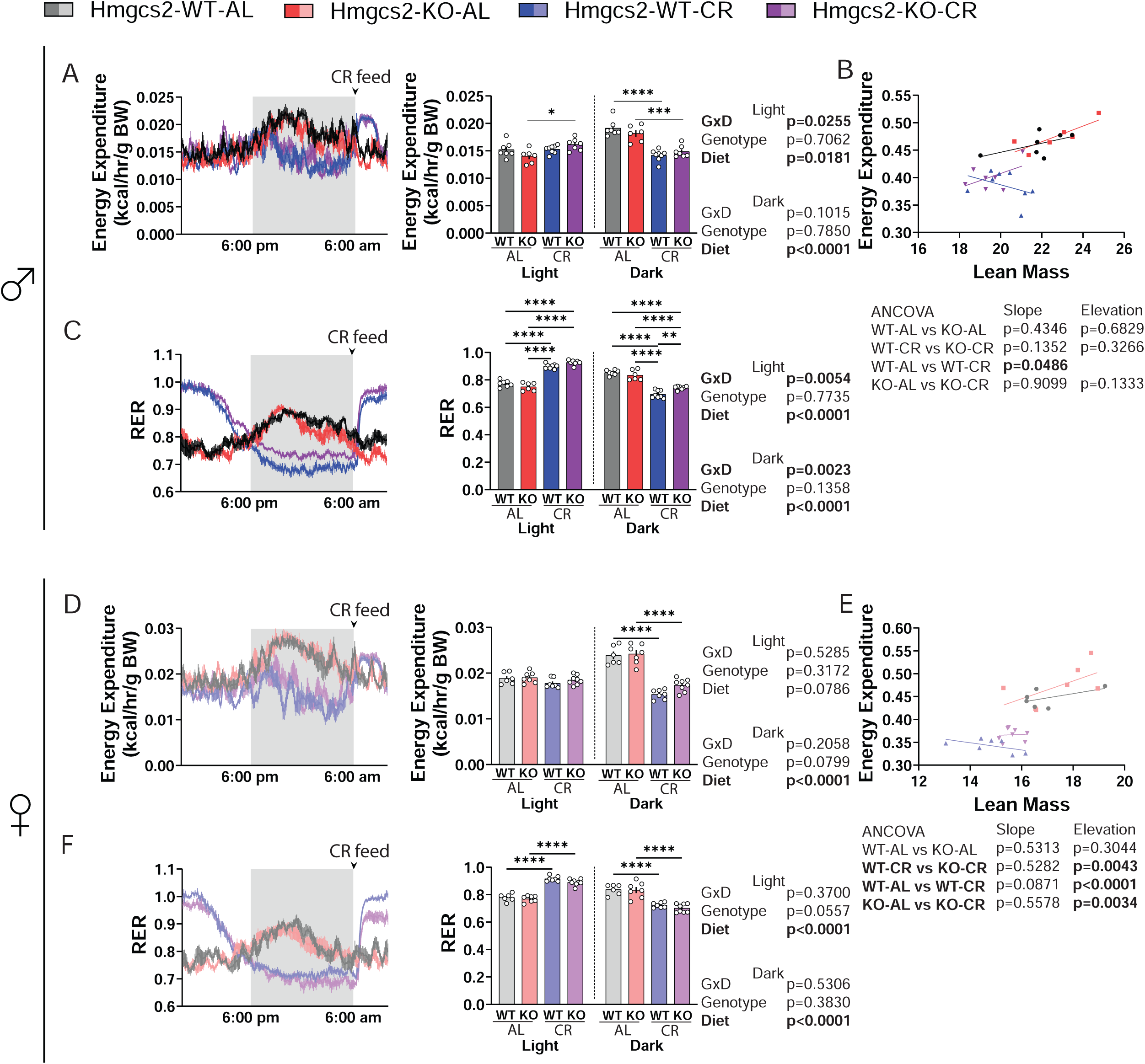
The ablation of ketogenesis induce sex-specific changes in RER. **(A-C)** Male mice energy expenditure (A) and respiratory exchange ratio (B) expressed over a 24 hrs period, binned by averaging all data points in the light or dark cycle. (C) ANCOVA analysis of average total energy expenditure against lean mass. For WT-AL, KO-AL, WT-CR, and KO-CR, n=7,6,8,7. **(D-F)** Female mice energy expenditure (D) and respiratory exchange ratio (E) expressed over a 24 hrs period, binned by averaging all data points in the light or dark cycle. (F) ANCOVA analysis of average total energy expenditure against lean mass. For WT-AL, KO-AL, WT-CR, and KO-CR, n=6,7,7,8. (A,C,D,F) *p<0.05, **p<0.01, ***p<0.001, ****p<0.0001, Sidak’s test post 2-way ANOVA, completed separately for each cycle. Data presented as mean ± SEM.

### *Hmgcs2* deletion selectively alters energy balance

We utilized indirect calorimetry to determine the effects of *Hmgcs2* knockout on CR metabolism. In the males, CR significantly increased energy expenditure of the KO mice in the light cycle and there was a genotype-diet interaction (**Fig. 3A**). In both males and females, CR significantly reduced energy expenditure in the dark cycle, reflecting a time when the animals would be fasting (**Figs. 3A & 3D**). To gain further insight on the animals’ energy expenditure, we performed ANCOVA analysis using lean mass as a covariate. The results confirmed that CR reduces overall energy expenditure in the females, and during the dark cycle in both sexes; interestingly, the loss of *Hmgcs2* slightly increased energy expenditure in female mice during CR (**Figs. 3B & 3E; Supplemental Figs. 4A-B & 4D-E**). The measurement of respiratory exchange ratio is an indicator of fuel selection. Due to the distinct feeding pattern of the CR animals, there was a significant, but expected, increase in RER during the light cycle and decrease in RER during the dark cycle of both male and female mice (**Figs. 3C & 3F**). In males, there was a genotype x diet effect in the light cycle and the dark cycle, the latter of which shows significantly increased RER in the KO mice treated with CR during the fasting period (**Fig. 3C**). In females, there was no significant effect of genotype on RER, although the post CR-feeding RER increase in the light cycle appears to be dampened by the loss of *Hmgcs2* (**Fig. 3F**). CR-feeding led to an increase in activity in the light cycle for both males and females; the average activity level was not different in any of the groups in the dark phase (**Supplemental Figs. 4C & 4F**).

### CR effects on iWAT adipocyte physiology and liver autophagy do not require *Hmgcs2*

We have previously shown that CR induces a significant reprogramming of adipocyte metabolism (Calubag et al., 2022). We characterized the inguinal white adipose tissues (iWAT) in WT and KO mice after 7 months of CR feeding. At the time of sacrifice, the total weight of iWAT was less in the CR animals (**Figs. 4B & 4G, Supplemental Figs. 5A & 5C**), and CR robustly reduced adipocyte size (**Figs. 4A, 4C, 4F, 4H**). The loss of *Hmgcs2* increased total iWAT mass in AL-fed males only (**Figs. 4B & 4G**). qPCR analysis of the iWAT of male mice treated with CR found robust upregulation of the thermogenic gene *Ucp1*, and genes that facilitate lipid metabolism, including *Dgat1*, *Elovl3*, *Cidea*, *Fasn*, and *Acc1*. Interestingly, the loss of *Hmgcs2* in males upregulated *Ucp1* even further and blunts the effect of CR on the expression of *Cidea* (**Figs. 4D-4E**). In the iWAT of the females, the same set of genes were likewise significantly upregulated by CR, except for *Ucp1* and *Acc1,* with no effect of *Hmgcs2* on the expression of *Ucp1* or *Cidea* (**Figs. 4I-4J**). In addition, the knockout of *Hmgcs2* suppressed *Lipe* and *Atgl* in females, but not in males (**Supplemental Figs. 5B & 5D**).

**Figure 4.**
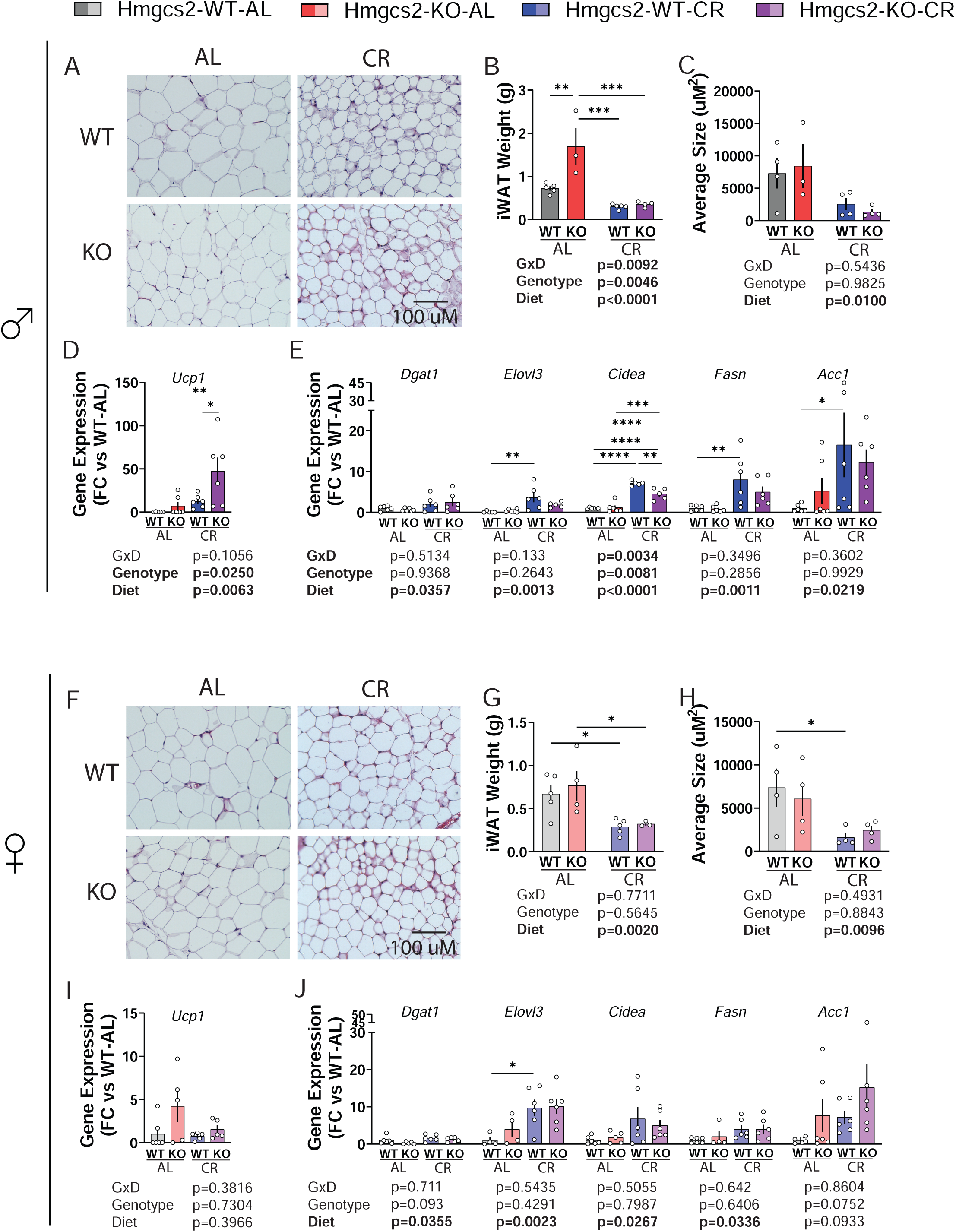
Loss of ketogenesis does not alter the effects of CR on iWAT physiology. **(A-C)** In male mice, representative H&E staining of the iWAT (A), iWAT weight (B) and the average size of adipocytes (C). For WT-AL, KO-AL, WT-CR, and KO-CR, iWAT weight n=5,3,5,4; adipocyte size n=4,3,4,4. **(D-E)** Male iWAT gene expression of *Ucp1* (D), n=6 each group; iWAT gene expression level of lipid processing genes (E) *Dgat1*, *Elovl3*, *Cidea*, *Fasn*, and *Acc1*. n=6 each group for *Dgat*1, *Fasn*, and *Acc1*; for WT-AL, KO-AL, WT-CR, and KO-CR, Elovl3 n=5,6,6,5; *Cidea* n=6,5,6,5. **(F-H)** In female mice, representative H&E staining of the iWAT (F), iWAT weight (G) and the average size of adipocytes (H). For WT-AL, KO-AL, WT-CR, and KO-CR, iWAT weight n=5,4,5,3; adipocyte size n=4 each group. **(I-J)** Female iWAT gene expression of *Ucp1* (I). For WT-AL, KO-AL, WT-CR, and KO-CR, n=6,5,6,6. Female iWAT gene expression level of lipid processing genes (J) *Dgat1*, *Elovl3*, *Cidea*, *Fasn*, and *Acc1*. n=6 each group for *Dgat1* and *Acc1*; *Elovl3* n=4,6,4,6; *Cidea* n=6,6,5,6; Fasn n=6,6,4,6. (B-E & G-J) *p<0.05, **p<0.01, ***p<0.001, ****p<0.0001, Sidak’s test post 2-way ANOVA (E & J) conducted separately for each gene. Data presented as mean ± SEM.

Autophagy is a critical function for healthy aging and has been linked to many lifespan-extending treatments (Kaushik et al., 2021). Autophagy is upregulated by CR and can also be induced by either a ketogenic diet (Liskiewicz et al., 2021) or βHB signaling (Abdelhady et al., 2023; Gomora-Garcia et al., 2023; Roohy et al., 2024). In the male liver, CR robustly upregulated the total expression of the autophagosome marker LC3 and the autophagy receptor p62 regardless of genotype (**Fig. 5A**). On the other hand, LC3 B/A ratio, and mTORC1-associated protein phosphorylation on pS757-ULK, pS240/244-S6, and pT37/46-4E-BP1 were unchanged (**Supplemental Fig. 5E**). In females, we did not detect any changes in the expression of autophagy proteins (**Fig. 5B**), but there was a slight decrease in the phosphorylation of pT37/46-4E-BP1 induced by CR (**Supplemental Fig. 5F**). There was no effect of genotype or a genotype-diet interaction on any of the protein targets we investigated.

**Figure 5.**
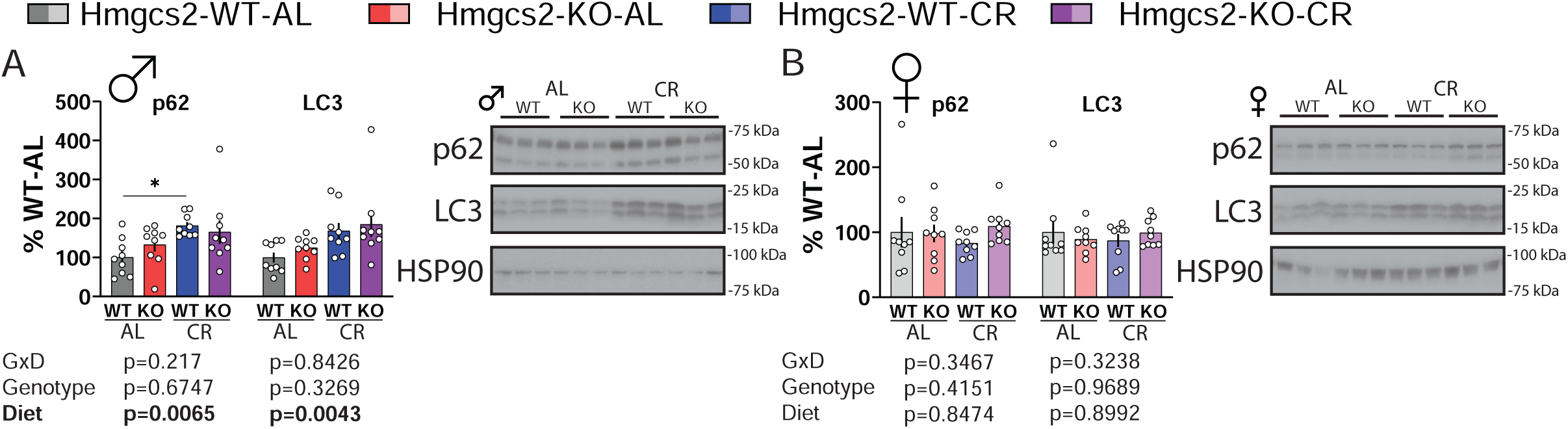
Caloric restriction upregulates autophagy-related proteins in the liver regardless of ketogenesis. (A-B) Liver western blots in male (A) and female (B) mice for p62 and LC3. n=9 each group. *p<0.05, Sidak’s test post 2-way ANOVA conducted separately for each protein. Data presented as mean ± SEM.

### Ketogenesis engagement is minimized after prolonged CR

To further understand why the loss of *Hmgcs2* had no overt effects on CR physiology, we sought to determine the ketone levels of these animals throughout the course of a day. We measured βHB over a period of 24 hrs, with each group in their typical experimental condition (**Fig. 6A; Supplemental Figs. 6A-6B**). Although WT mice on the CR diet exhibited slightly higher βHB levels than those on the AL diet, the differences were negligible at most timepoints, with peak βHB concentrations remaining below 1.0 mmol/L. This contradicts the long-held assumption that CR should significantly increase ketone body production (de Cabo & Mattson, 2019). To further investigate this phenomenon, we measured the circulating βHB level of the AL mice while fasting, starting at the beginning of the light cycle (**Fig. 6B; Supplemental Figs. 6C-6D**). As expected, ketone levels raised in the WT group only, with ketogenesis particularly engaged after the dark cycle begins. After 12-16 hrs of fasting, circulating βHB levels peaked at 2.5+ mmol/L, much higher than that achieved by the mice treated with CR. For comparison, we placed a dotted line at the 1 mmol/L level in both graphs. This rise in βHB level was not observed in the fasting AL-KO group, which still exhibited fluctuations in βHB readings, but at a much lower level.

**Figure 6.**
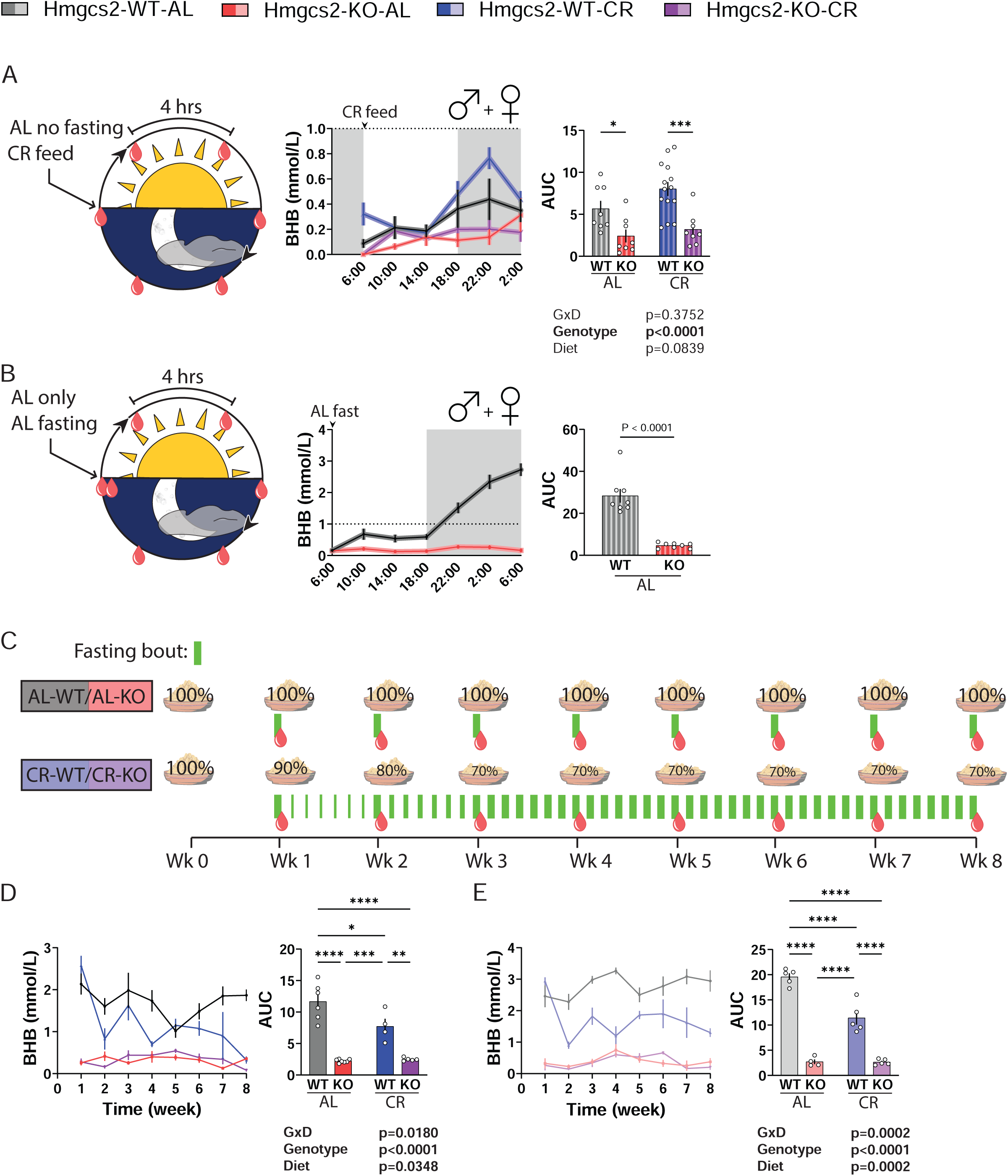
Acclimation to a recurrent fasting caloric restriction protocol suppresses ketogenesis. **(A)** Circulating βHB level of male + female mice measured every 4 hrs under typical experimental conditions (food present for AL groups/CR groups fed at 6 am). For WT-AL, KO-AL, WT-CR, and KO-CR, n=8,8,14,8. **(B)** Circulating βHB level of typically *ad libitum*-fed male + female mice fasting starting at 6 am. n=8 each group. **(C)** Diagram of the CR acclimation experiment using a separate cohort of mice. **(D-E)** Fasting βHB level of male (D) and female (E) mice during the acclimation phase of the CR protocol. For WT-AL, KO-AL, WT-CR, and KO-CR, male n=6,7,4,5; female n=5,4,5,5. (A & D-E) *p<0.05, **p<0.01, ***p<0.001, ****p<0.0001, Sidak’s test post 2-way ANOVA. (B) *p<0.05, t-test. Data presented as mean ± SEM.

Finally, we looked to further characterize the development of this resistance to ketogenesis by CR-fed mice by measuring fasting ketone levels during the first 8 weeks of CR initiation. In a separate cohort of mice, we measured the fasting ketone levels of all groups once per week at the conclusion of a 21 hrs fast. As part of this experiment, AL-fed mice were fasted only once per week, whereas CR-fed mice experienced self-administered daily fasting as a result of their feeding pattern (**Fig. 6C**). We found that in both males and females, the circulating βHB level achieved by the WT CR-fed mice begin to drop below that of the WT AL-fed mice after 2 weeks of CR conditioning (**Figs. 6D-6E**). These results are evidence that the adaptation to daily fasting in CR-fed mice results in drastically reduced circulating ketone levels in response to fasting.

### Caloric restriction enhances ketone body metabolism

This significant reduction in circulating ketone levels could result from either a lack of ketone body production or an increase in ketone body consumption. To determine whether the ability of these animals for ketogenesis and ketolysis is altered, we treated them with either exogenous medium-chained triglycerides (MCT) or βHB, respectively. Consumption of βHB-precursors, such as medium triglyceride (MCT) oil, acutely increases circulating βHB in humans (Heidt et al., 2023; Norgren et al., 2020) and in mice (Shoji, Kunugi, & Miyakawa, 2022). After oral administration of MCT oil, we monitored the circulating level of βHB for 4 hrs. We found that this treatment reliably engaged ketogenesis, particularly in females, with WT female mice fed the CR diet showing significantly higher levels of circulating βHB than AL-fed WT females (**Figs. 7A-7B**).

**Figure 7.**
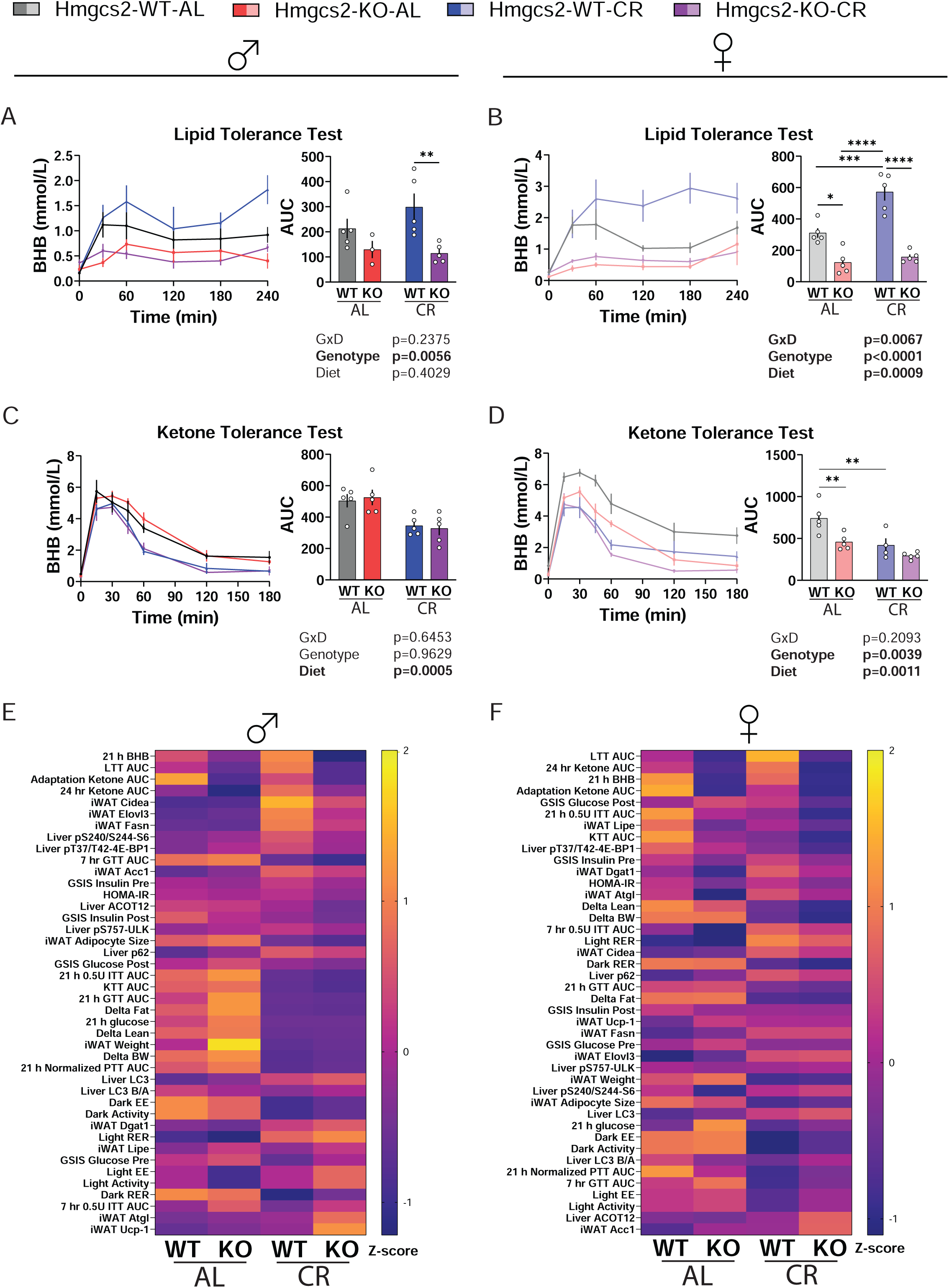
Caloric restriction enhances ketogenesis and ketolysis capacity. **(A-B)** Lipid tolerance test using MCT oil (5 ml/kg; P. O.) in male (A) and female (B) mice. Circulating βHB is unchanged in the KO mice of both sexes while CR significantly enhances ketogenic ability in the lipid tolerance test. For WT-AL, KO-AL, WT-CR, and KO-CR, male n=5,3,5,5; female n=5 each group. **(C-D)** Ketone tolerance test using sodium 3-β-hydroxybutyrate in saline (2 g/kg; I. P.) 4 hrs after CR-feeding and AL fasting. Ketone consumption is improved in male (C) and female (D) mice treated with CR. Male n=5 each group; for WT-AL, KO-AL, WT-CR, and KO-CR, female n= 5,5,4,5. **(E-F)** Heatmap representation of the discrete dataset in this study for male (E) and female (F), expressed as Z-score comparisons within individual variables, sorted by the difference of WT-CR vs KO-CR. (A-D) *p<0.05, **p<0.01, ***p<0.001, ****p<0.0001, Sidak’s test post 2-way ANOVA. Data presented as mean ± SEM.

In the ketone tolerance test, we injected exogenous sodium βHB and monitored circulating βHB levels for 3 hrs. This approach has been shown to readily raise circulating βHB levels in mice (Kraeuter, Mashavave, Suvarna, van den Buuse, & Sarnyai, 2020; Sun et al., 2022; Yum, Ko, & Kim, 2012). We confirmed a rapid absorption of the exogenous βHB into circulation, where the peak level was observed 15-30 min after injection and steadily decreased thereafter. In both males and females, CR feeding clearly enhanced the metabolism of this excess ketone challenge (**Figs. 7C-7D**). In females only, the loss of *Hmgcs2* further enhanced ketone body catabolism, significantly decreasing the βHB area under the curve in AL-fed mice. Together, these results suggest that CR mice have enhanced capacity for both ketogenesis and ketolysis if the appropriate substrates are available.

Finally, recent work provided evidence that acetate production may be an alternate pathway for energy production from fatty acids when ketogenesis is suppressed (Feola et al., 2024). This increase in circulating acetate is incredibly transient, therefore we attempted to look at upstream markers of this adaptation. The conversion of acetyl-CoA to acetate in the liver is regulated by acetyl-coenzyme A thioesterase 12 (ACOT12), which is upregulated in fasted mice lacking HMGCS2 (Feola et al., 2024) and has been shown to regulate the expression of ketogenic genes, including *Hmgcs2* (Wang et al., 2023). To understand whether ACOT12 protein expression is altered in CR and by the loss of *Hmgcs2*, we probed for ACOT12 and found that there was a small but significant increase in the hepatic ACOT12 of the female CR mice (**Supplemental Figs. 6E-6F**). This suggests that acetate may partially compensate for ketone production during CR.

To summarize our findings, a visualization of all discrete data is rank-sorted by the difference between the WT and KO mice under the CR feeding condition and provided in heatmap form (**Figs. 7E-7F**). As expected, the top traits involve ketone body metabolism, while the arrangement of the remaining data was less predictable.

## Discussion

The precise mechanisms by which geroprotective dietary interventions promote healthy aging continue to be elusive. Here, we provided clear evidence that ketogenesis is dispensable for the many metabolic benefits of a CR diet, and that adaptation to daily fasting simply discourages βHB production. Nevertheless, the current literature has conflicting reports on whether the CR diet actually increases circulating βHB. In mice fed a 40% CR diet once per day, fasting blood βHB levels appeared to decrease, with a counterbalanced enhancement in liver glycogen (Hu et al., 2022). To this aim, the contribution of enhanced glycogen metabolism on physiological health has yet to be elucidated. In aged rats, a 40% CR diet fed once per day increased circulating ketone levels (Lin, Zhang, Gao, & Watts, 2015). In a mouse model of prostate cancer, a two-days-a-week (5:2) intermittent fasting protocol caused urinary ketone to be initially highly elevated, but returned to baseline levels after 8 fasting episodes (J. A. Thomas, 2nd et al., 2010).

Our ability to interpret human data to determine if this response is conserved is limited, as most published studies have either a low number of participants or lack a control group. Several studies have shown that fasting or CR in overweight or obese subjects do induce ketogenesis. In a patient case report, a 5:2 fasting protocol with 36-hrs fasting windows observed that circulating βHB remained highly elevated even after 8 months of practice (Borer, 2024). Obese (BMI >30) asthma patients treated with alternate day 20% CR reliably have increased ketone production for at least 8 weeks (Johnson et al., 2007). However, some very long-term studies of CR suggest that our findings also apply to people. In a study of young women (BMI 24-40) treated with 25% daily energy restriction, there was an initial increase in serum ketone level at month 1 and 3, but this difference was reduced with time and disappeared by month 6; meanwhile in the same study, 25% CR in the form of severe restriction for 2 day/week appears to maintain elevated ketone levels over 6 months (Harvie et al., 2011). The phase II CALERIE clinical trial was one of the most comprehensive CR patient studies, and unique in that it focused on non-obese adults. After 24 months, patients practicing ∼11.9% CR (prescribed as 25%) exhibited a modest increase in fasting plasma ketone bodies compared to the AL group (p=0.055; (Huffman et al., 2022)), with a very low incidence of urinary ketosis (4.4% in CR vs 1.9% in AL; (Ferguson, Sahoo, McGrail, Francois, & Stratton, 2022)). On balance, these studies suggest that while CR and fasting initially induce ketogenesis in the overweight and obese, over the long-term CR does not drastically elevate ketone levels in most individuals.

In addition to its role as an energy currency, βHB is known to act as an effective signaling molecule that elicits multiple benefits (Newman & Verdin, 2017). βHB is an inhibitor of histone deacetylases (HDACs) (Shimazu et al., 2013) and contributes to a unique post-translational protein modification, lysine β-hydroxybutyrylation (Huang et al., 2021). While the concentration of βHB that needs to be reached for its signaling properties has not been established, our results suggest that the βHB levels reached during CR may be a poor surrogate to investigate this, as βHB concentration in a CR-treated mouse is not higher than that of a non-fasting AL animal. This implies that many CR benefits are achieved either through another mechanism distinct from the actions of βHB, or that the effect of βHB only requires a small local increase that is undetectable in circulation.

Acetoacetate is the established precursor to βHB. While the conversion between the two molecules is reversible, the deletion of hepatic *Hmgcs2* has been shown to deplete both plasma acetoacetate and βHB (Feola et al., 2024). However, despite the deletion of *Hmgcs2* in our study, we observed non-zero amounts of circulating βHB. This has been previously observed by other groups (Feola et al., 2024), and is most likely explained by two known routes of non-canonical ketogenesis. First, endogenous gut microbiota can produce butyrate, which can directly contribute to plasma βHB levels through a molecular pathway independent of HMGCS2 (Chen et al., 2023). Further, the branched chain amino acid leucine can be catabolized to produce HMG-CoA, the precursor to acetoacetate, without HMGCS2, accounting for roughly 4.4% of the ketone bodies in starved rats (L. K. Thomas, Ittmann, & Cooper, 1982). Nevertheless, it is clear from our data that these alternate sources of ketone body are dwarfed by canonical *Hmgcs2*-dependent ketogenesis, which is completely abolished by the deletion of *Hmgcs2* and not compensated for by the non-canonical pathways. In addition, acetate production from acetyl-CoA has been proposed as an alternative pathway for metabolizing fatty acids when ketogenesis is ablated (Feola et al., 2024). Circulating acetate and the expression of *Acot12* were observed to be higher after 24 hrs fasting but are unchanged in the fed state (Feola et al., 2024). It is likely that we only observed a small increase in ACOT12 protein in CR-fed animals as we collected tissues only a few hours after feeding.

We observed several small but significant physiological adaptations with the loss of HMGCS2. First, female KO mice, regardless of whether they were fed an AL or CR diet, have a stronger response to insulin, as well as a trend towards an improved HOMA-IR. There is a robust and sex-specific adaptation in fuel selection, where the absence of HMGCS2 during CR increased RER in the males and decreased RER in the females; KO females also had increased energy expenditure during CR. Changes in liver gluconeogenesis and glycogen storage in mice lacking HMGCS2 or treated with CR has been demonstrated previously (d’Avignon et al., 2018; Hu et al., 2022). Future work will need to be conducted to understand how these adaptations contribute to these physiological responses we observed. In addition, recent reports found mice lacking HMGCS2 to exhibit a rapidly reversible fatty liver and hepatic steatosis phenotype when challenged (Queathem et al., 2025); however, we did not observe any instances of fatty liver in our KO mice, perhaps due to the transient nature of these conditions and the timing of our tissue collection.

Another limitation in this study is that we cannot test the role of HMGCS2 in every CR response, owing to the incredibly wide range of effects induced by CR. While we focused on metabolism, CR can elicit other effects that demand their own investigation, including improved brain function (Hadem, Majaw, Kharbuli, & Sharma, 2019), mitochondria health (Lanza et al., 2012), and disease mitigation (Green et al., 2022). Similarly, we only evaluated these effects in one strain of mouse and did not evaluate the role of HMGCS2 in CR-induced lifespan extension. The role of ketone bodies in different genetic background, and its roles on lifespan is an important future study, as there have been mouse strains identified for which CR do not extend or even shorten lifespan (Liao, Rikke, Johnson, Diaz, & Nelson, 2010; Mitchell et al., 2016; Mulvey et al., 2021). Interestingly, *Hmgcs2* has been lost during the evolution of several long-lived mammals, such as whales and elephants, suggesting that further optimizations of ketone body metabolism may augment mammalian longevity (Jebb & Hiller, 2018).

Lastly, there are many permutations of the CR protocol. Here, we focused solely on one of the most traditional versions of CR, with feeding carried out once daily at the beginning of the light cycle. We have previously shown that the metabolic phenotypes, especially insulin sensitivity, are largely preserved regardless of the feeding time (Pak et al., 2024), and while some studies suggest time of feeding impacts the degree to which CR extends lifespan, it is clear that CR with once-daily feeding at the beginning of the light cycle robustly extends lifespan (V. Acosta-Rodriguez et al., 2022; Nelson & Halberg, 1986; Pak et al., 2021). Since ketogenesis is diametrically opposed by insulin, and is directly inhibited by insulin signaling (Kolb et al., 2021), we believe our main findings are likely robust to differences in time of feeding.

It is a long-held assumption that the fasting component of CR induces significant ketogenesis, contributing to the benefits of the diet (de Cabo & Mattson, 2019). Here, we provided proof that this assumption – like many others about the mechanisms by which CR functions (Pearson et al., 2008; Yu et al., 2019) – may be incorrect. Mice completely lacking the capacity for fasting-induced ketogenesis still receive most of the metabolic benefits of CR, and indeed, long-term adaptation to CR downregulates ketogenesis. Further work should be done to understand the mechanisms of this physiological “memory” for fasting. Altogether, our result demonstrates that it may be wise for us to consider the effects of dietary interventions not solely as acute responses, but more as part of a biological framework consisting of long-term adaptations.

## Materials and Methods

### Statistical analysis

Where an analysis for a 2-factorial experimental design is possible, we performed 2-way ANOVA with Sidak’s multiple comparison post hoc test when the main effects or the interaction are significant. Exact p-value of the post hoc tests are provided in the source data file. All significance test calculations were done with GraphPad Prism ver10.4.0.

### Animals

All procedures were performed in accordance to institutional guidelines and were approved by the Institutional Animal Care and Use Committee of William S. Middleton Memorial Veterans Hospital (Madison, WI, USA) and the University of Wisconsin-Madison. *Hmgcs2* KO mice were generated by the Advanced Genome Editing Lab of the University of Wisconsin-Madison Biotechnology Center, which performed pronuclear microinjection of a CRISPR/Cas9 construct that resulted in the insertion of a fragmented gene duplication downstream of exon 2, rendering the loss of functional gene transcript expression. This was confirmed with long-read (PacBio) genome sequencing. The genetic background of the original animal was C57BL/6J. *Hmgcs2* heterozygotes were bred to generate matched WT and KO lines, which were maintained separately. The loss of detectable HMGCS2 protein and the ability to engage ketogenesis was confirmed as presented.

All animals in this experiment were singly housed with a plastic hut and nesting materials. Calorie-free aspen bedding materials were changed weekly. Food consumption of the AL groups was monitored weekly and the values used to adjust the CR food allotment for the following week. CR-fed mice were fed daily at the beginning of the light phase, typically between 6:00 to 7:00 am. All mice were fed the Teklad Global 18% Protein Rodent Diet (Envigo 2018). CR was implemented in a step-down protocol of increasing 10% per week until 30% restriction is reached. Animals were adapted to the feeding protocol for 8 weeks prior to physiological testing. A schematic of this workflow is presented in **Supplemental Fig. 1A**. All mice were maintained at a temperature of 22⁰C, and health checks were performed daily by the facility staff. Relative humidity of the animal room ranges between 30-70%, with an average of 42%. Mice were housed in a SPF facility in static microisolator cages under 12:12 light cycle conditions with *ad libitum* access to water. Body composition was determined using an EchoMRI Body Composition Analyzer.

Animals were temporarily housed in the Oxymax/CLAMS-HC metabolic chamber system (Columbus Instruments) during indirect calorimetry experiments. Mice were housed in this system for ∼48 continuous hours. The first ∼24 hours of data was discarded as acclimation period with a subsequent continuous 24 hours period utilized for data analysis.

### RT-qPCR

CR mice were euthanized 4-5 hrs after feeding via cervical dislocation and AL mice were fasted overnight and refed at the same time as CR feeding. Tissues were flash frozen using L_2_N. mRNA extraction was performed on iWAT using TRI reagent (AM9738; Invitrogen) and cDNA was generated using Superscript III reverse transcriptase (56575; Invitrogen) according to manufacturer instructions. RT-qPCR was carried out using SYBR green (*Power* SYBR Green PCR Master Mix; 4367659; Applied Biosystems). The following primers were used: *Dgat1* F 5’TGG TGT GTG GTG ATG CTG ATC’3 R 5’GCC AGG CGC TTC TCA A3’; *Elovl3* F 5’ATG CAA CCC TAT GAC TTC GAG 3’ R 5’ACG ATG AGC AAC AGA TAG ACG 3’; *Cidea* F 5’GAA TAG CCA GAG TCA CCT TCG 3’, R 5’AGC AGA TTC CTT AAC ACG GC3’; *Fasn* F 5’CCCCTCTGTTAATTGGCTCC 3’, R 5’TTGTGGAAGTGCAGGTTAGG 3’; *Acc1* F 5’AAG GCT ATG TGA AGG ATG 3’, R 5’CTG TCT GAA GAG GTT AGG 3’; *Ucp1* F 5’ GCA TTC AGA GGC AAA TCA GC 3’, R 5’ GCC ACA CCT CCA GTC ATT AAG 3’; *Lipe* F 5’ CTG AGA TTG AGG TGC TGT CG 3’, R 5’ CAA GGG AGG TGA GAT GGT AAC 3’; *Atgl* F 5’ ATA TCC CAC TTT AGC TCC AAG G 3’, R 5’ CAA GTT GTC TGA AAT GCC GC 3’; *β-Act* F 5’ GAT GTA TGA AGG CTT TGG TC 3’, R 5’ TGT GCA CTT TTA TTG GTC TC 3’. qPCR outlier analysis was performed using Prism 10.4.0 built-in function ROUT (Q=1%).

### Western Blot

Liver tissues for Western Blotting were flash frozen using L_2_N and homogenized in RIPA buffer containing EDTA-free Protease and Phosphotase Inhibitor Mini Tablet (Thermo Scientific, A32961). Quantification of all proteins were normalized to the loading control and expressed as a percentage of the WT-AL group. Total protein quantifications were normalized to HSP90 and phosphoresidues were normalized to the total protein. Antibodies were purchased from Cell Signaling Technology: Hmgcs2 (20940), LC3A/B (12741), HSP90 (4877), ULK1 (8054), pS757-ULK (14202), 4E-BP1 (9644), pT37/46-p4E-BP1 (2855), S6 (2217), pS240/244-S6 (2215), HRP-linked anti-rabbity secondary (7074). The P62 antibody was purchased from Progene Biotechnik (GP62-C). The ACOT12 antibody was purchased from Sigma-Aldrich (SAB2100029). HRP-linked anti-guinea pig secondary antibody was purchased from American Research Products (90001).

### *In vivo* procedures

Metabolic tolerance tests were carried out after either 21 hrs or 7 hrs fasting as indicated. The 21 hrs fast tests were carried out at the beginning of the light cycle while the 7 hrs fast tests were carried out ∼3 hrs prior to the dark cycle. In the glucose and insulin tolerance test, mice were injected with glucose (1 g/kg; Sigma-Adrich, G7021) or insulin (0.50 U/kg; Novolin) intraperitoneally (I.P.). For glucose-stimulated insulin secretion, 2 g/kg of glucose was used. Blood glucose was monitored via a Bayer Contour glucometer. Lipid tolerance test (LTT) and ketone tolerance test (KTT) were performed on animals after ∼4 hrs fast post the CR feeding period. In the KTT, βHB in the form of a sodium salt (CAS-No: 150-83-4; Sigma-Aldrich) is administered at 2 g/kg via I.P. injections. In the lipid tolerance test, MCT oil, containing mostly capric acid and caprylic acid (M4200; CAS 73398-61-5; Spectrum Chemical), was delivered via oral gavage (5 mL/kg). Ketone level was measured using KetoBM handheld ketone meter, which measures βHB through the activities of β-hydroxybutyrate dehydrogenase.

### iWAT Histology

After sacrifice, a portion of the iWAT was fixed in formaldehyde. Paraffin sections were stained with H&E and quantified using imageJ for adipocyte size. For each mouse, 2-4 fields of view were scored by three blinded researchers and averaged together for analysis.

## DATA AVAILABILITY STATEMENT

The data that support the plots within this article and other findings of this study, including full scans of western blot images, are provided as Source Data files.

## ACKNOWLEDGEMENTS

We would like to thank all members of the Lamming lab for their assistance and input. The Lamming laboratory is supported in part by the NIH/NIA (AG056771, AG081482, AG084156, AG085898 and AG094153 to D.W.L.), NIH/NIDDK (DK125859 to D.W.L.) and startup funds from the University of Wisconsin-Madison School of Medicine and Public Health and Department of Medicine to D.W.L. C-Y.Y. was supported in part by a NIA F32 postdoctoral fellowship (F32AG077916), a NIA K99/R00 award (K99AG084921), and a Research Enrichment Component Postdoctoral Scholarship from the Wisconsin Alzheimer’s Disease Research Center (P30-AG062715). M.F.C. is supported by F31AG082504. R.B. was supported by F31AG081115. M.E.T. is supported by a F99/K00 award AG083290. The authors used the University of Wisconsin-Madison Genome Editing and Animal Models (GEAM) Core for generating the animals and the UW Carbone Cancer Center Experimental Animal Pathology Laboratory (P30CA014520) for the iWAT staining. D.W.L. is a member of the Wisconsin Nathan Shock Center of Excellence in the Basic Biology of Aging, P30 AG092586. The Lamming lab was supported in part by the U.S. Department of Veterans Affairs (IS1-BX005524), and this work was supported using facilities and resources from the William S. Middleton Memorial Veterans Hospital. The content is solely the responsibility of the authors and does not necessarily represent the official views of the NIH. This work does not represent the views of the Department of Veterans Affairs or the United States Government.

