## Supplemental Figures for "Ketogenesis is dispensable for the metabolic adaptations to caloric restriction"

Supplemental Figure 1.

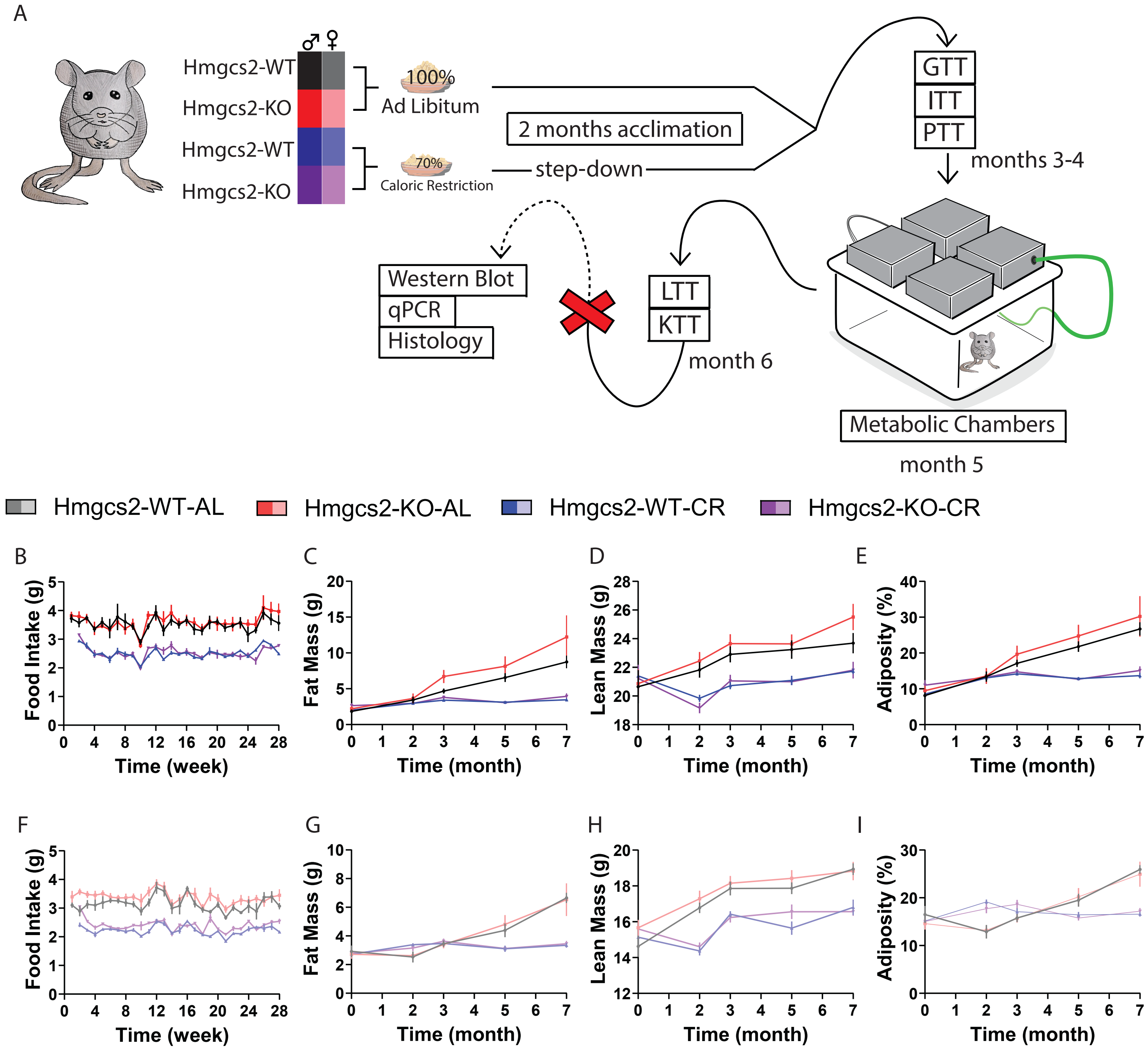

**Supplemental Figure Legends**

**Supplemental Figure 1. The ablation of ketogenesis does not affect the fat, lean, and adiposity of mice undergoing caloric restriction.**

**(A)** A schematic of our workflow. Abbreviations: GTT (glucose tolerance test), ITT (insulin tolerance test), PTT (pyruvate tolerance test), LTT (lipid tolerance test), and KTT (ketone tolerance test). **(B-E)** In male mice, their food intake (B), fat mass (C), lean mass (D), and adiposity % (E) over 7 months of the caloric restriction protocol. For all male mice data presented here, at the beginning of the experiment, WT-AL, KO-AL, WT-CR, KO-CR, n=7, 6, 8, and 7 respectively. **(F-I)** In female mice, their food intake (F), fat mass (G), lean mass (H), and adiposity % (I) over 7 months of the caloric restriction protocol. For all female mice data presented here, at the beginning of the experiment, WT-AL, KO-AL, WT-CR, KO-CR, n=6, 7, 7, 9 respectively. Data presented as mean ± SEM.

Supplemental Figure 2.

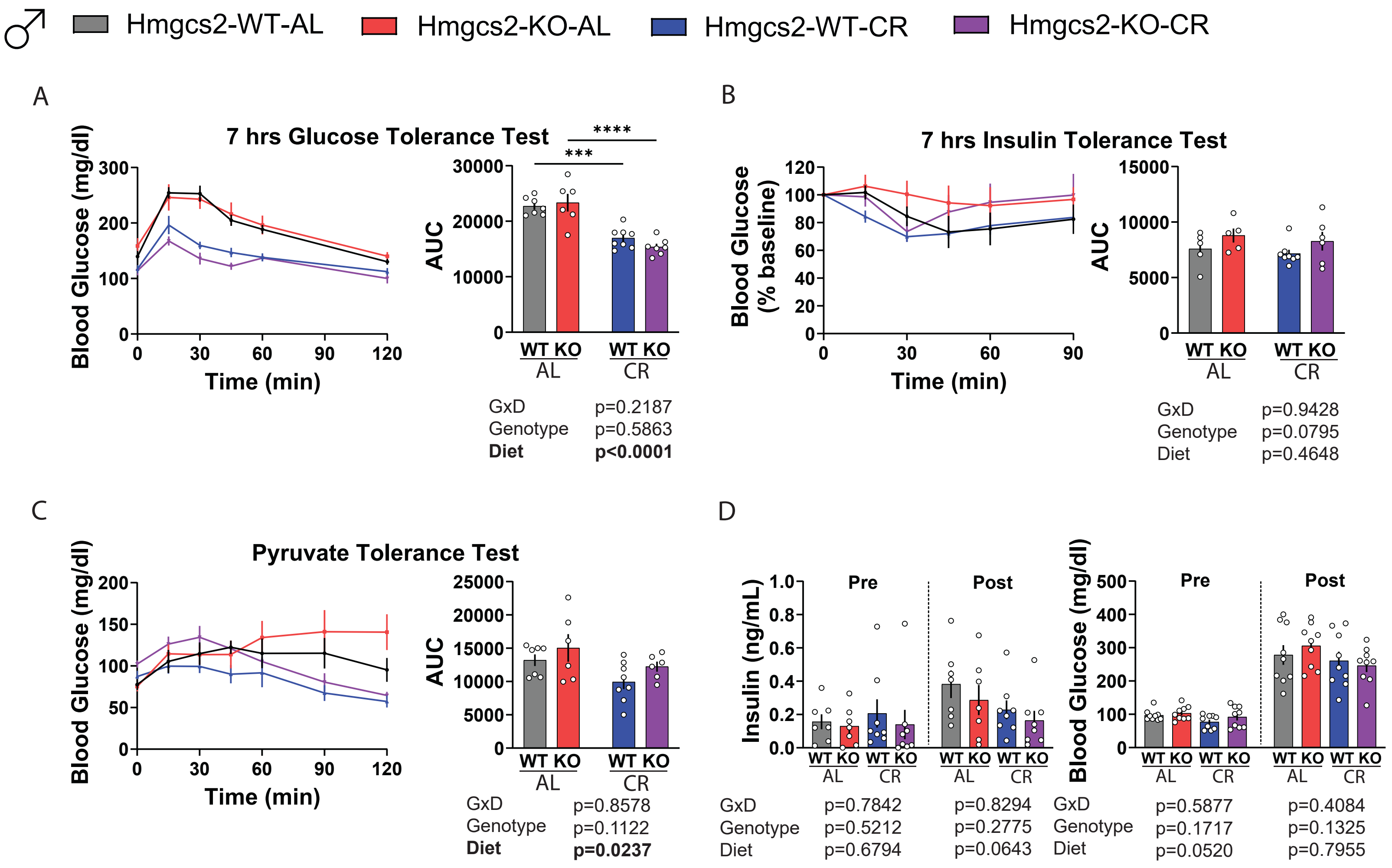

**Supplemental Figure 2. Male metabolic health characterization**

**(A-B)** After 7 hrs fast in male mice, glucose tolerance test (A; 1 g/kg; I. P.) and insulin tolerance test (B; 0.5 U/kg; I. P.). For WT-AL, KO-AL, WT-CR, and KO-CR, glucose tolerance test n=7,6,8,7; insulin tolerance test n=5,5,8,6. **(C)** In male mice, raw blood glucose value of the pyruvate tolerance test (2 g/kg; I. P.) after 21 hrs fasting. For WT-AL, KO-AL, WT-CR, and KO-CR, n=7,6,8,6. **(D)** After 21 hrs fasting in male mice, glucose stimulated insulin secretion (2 g/kg; I. P.) insulin and blood glucose levels. For WT-AL, KO-AL, WT-CR, and KO-CR, insulin n=7,7,8,8; glucose n=9 each group. (A-D) ***$p<0.001$, ****$p<0.0001$, Sidak's test post 2-way ANOVA. Data presented as mean ± SEM.

Supplemental Figure 3.

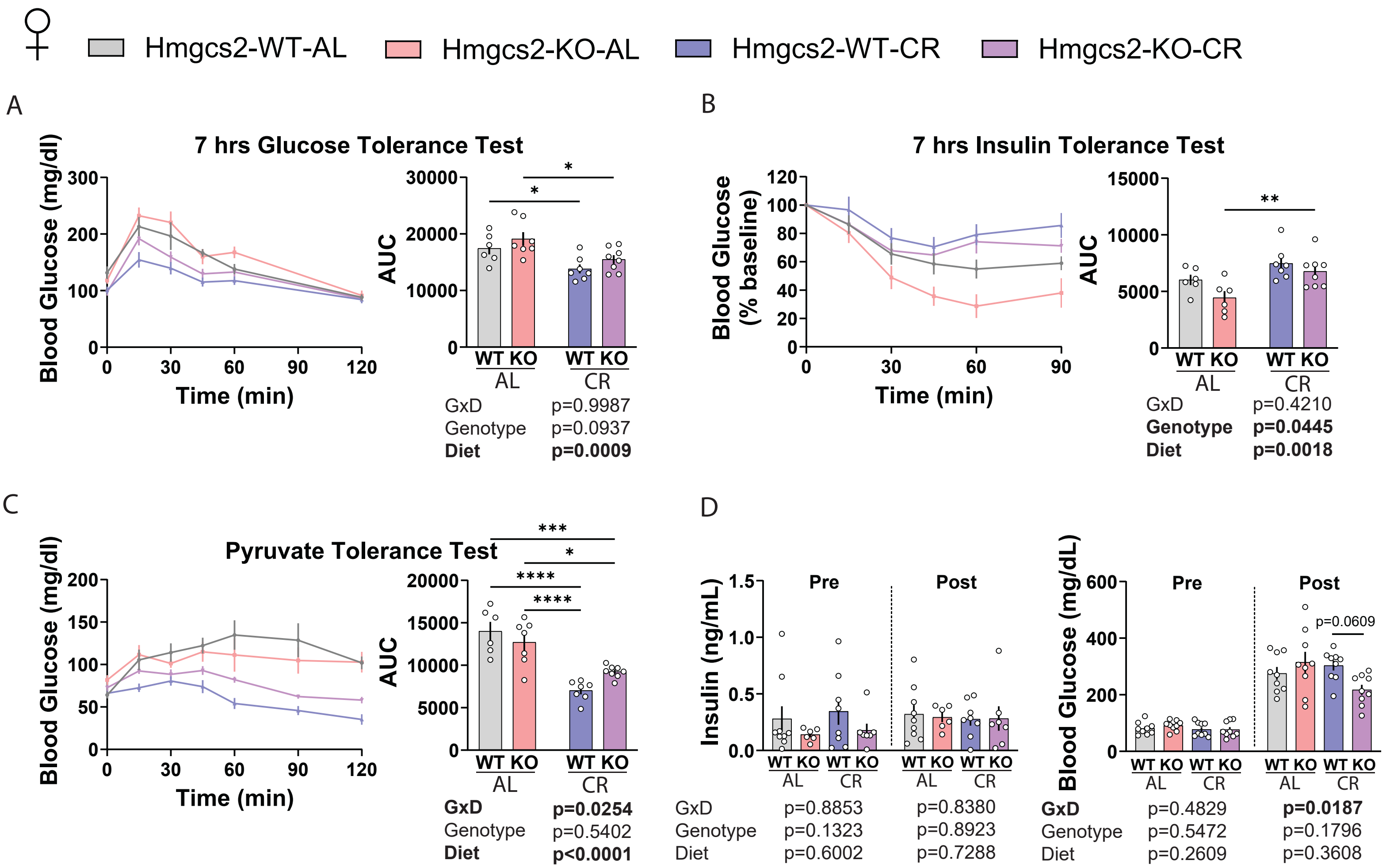

**Supplemental Figure 3. Female metabolic health characterization.**

**(A-B)** After 7 hrs fast in female mice, glucose tolerance test (A; 1 g/kg; I. P.) and insulin tolerance test (B; 0.5 U/kg; I. P.). For WT-AL, KO-AL, WT-CR, and KO-CR, glucose tolerance test n=6,7,7,8; insulin tolerance test n=6,6,7,8. **(C)** In female mice, raw blood glucose value of the pyruvate tolerance test (2 g/kg; I. P.) after 21 hrs fasting. For WT-AL, KO-AL, WT-CR, and KO-CR, n=6,7,7,8. **(D)** After 21 hrs fasting in female mice, glucose stimulated insulin secretion (2 g/kg; I. P.) insulin and blood glucose levels. For WT-AL, KO-AL, WT-CR, and KO-CR, insulin n=9,6,8,7; glucose n=9 each group. (A-D) *$p<0.05$, **$p<0.01$, ***$p<0.001$, ****$p<0.0001$, Sidak's test post 2-way ANOVA. Data presented as mean ± SEM.

Supplemental Figure 4.

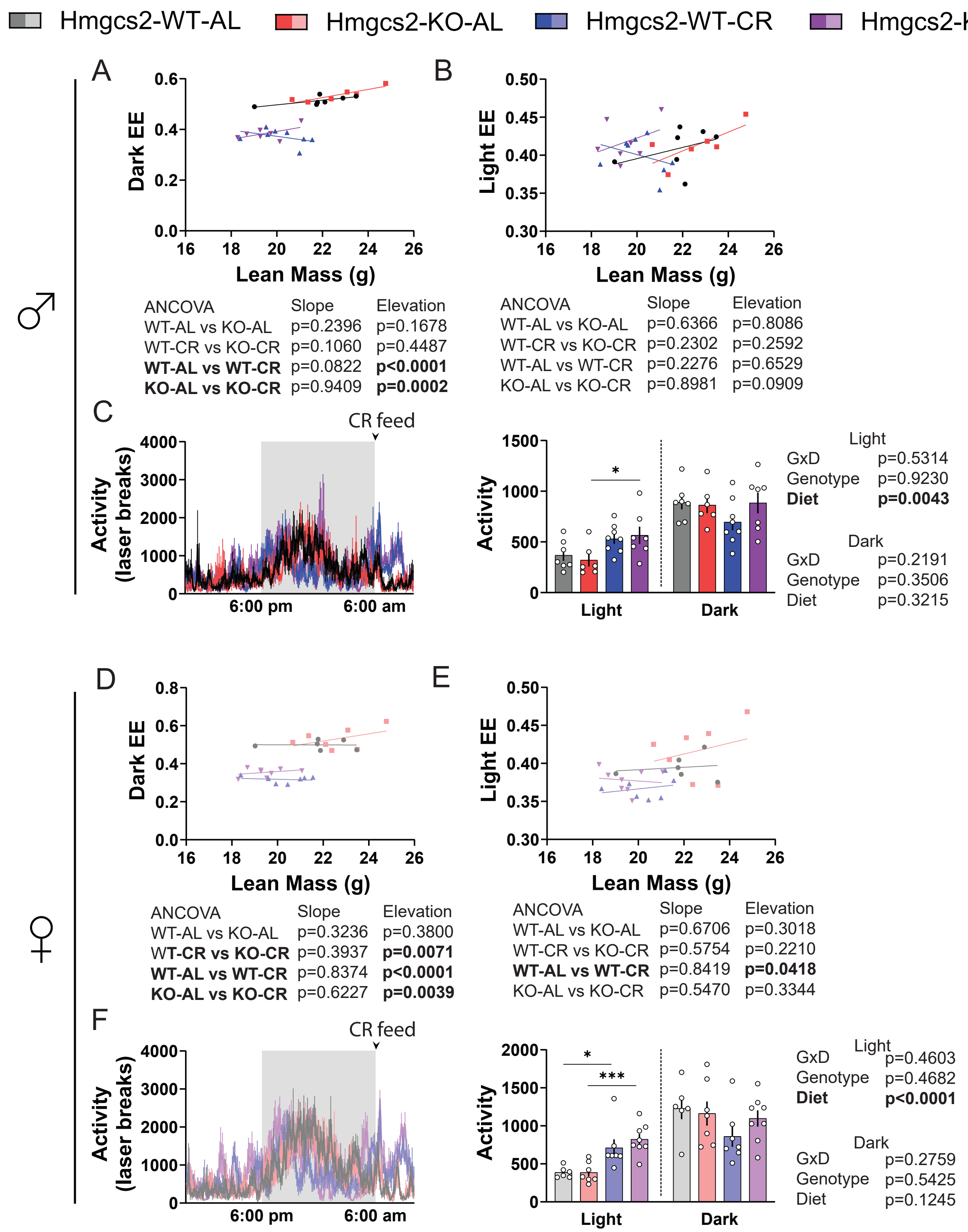

**Supplemental Figure 4. ANCOVA analysis and metabolic chamber activity.**

**(A-C)** In male mice, ANCOVA analysis of average dark phase (A) and light phase (B) energy expenditure against lean mass. (**C**) Activity during metabolic chambers test in male mice over 24 hrs and binned by averaging all data points in the light or dark cycle. For WT-AL, KO-AL, WT-CR, and KO-CR, male n=7,6,8,7. **(D-F)** In female mice, ANCOVA analysis of average dark phase (D) and light phase (E) energy expenditure against lean mass. (**F**) Activity during metabolic chambers test in female mice over 24 hrs and binned by averaging all data points in the light or dark cycle. For WT-AL, KO-AL, WT-CR, and KO-CR, female n=6,7,7,8. (C, F) *$p<0.05$, ***$p<0.001$; Sidak's test post 2-way ANOVA performed separately for each cycle. Data presented as mean ± SEM.

Supplemental Figure 5.

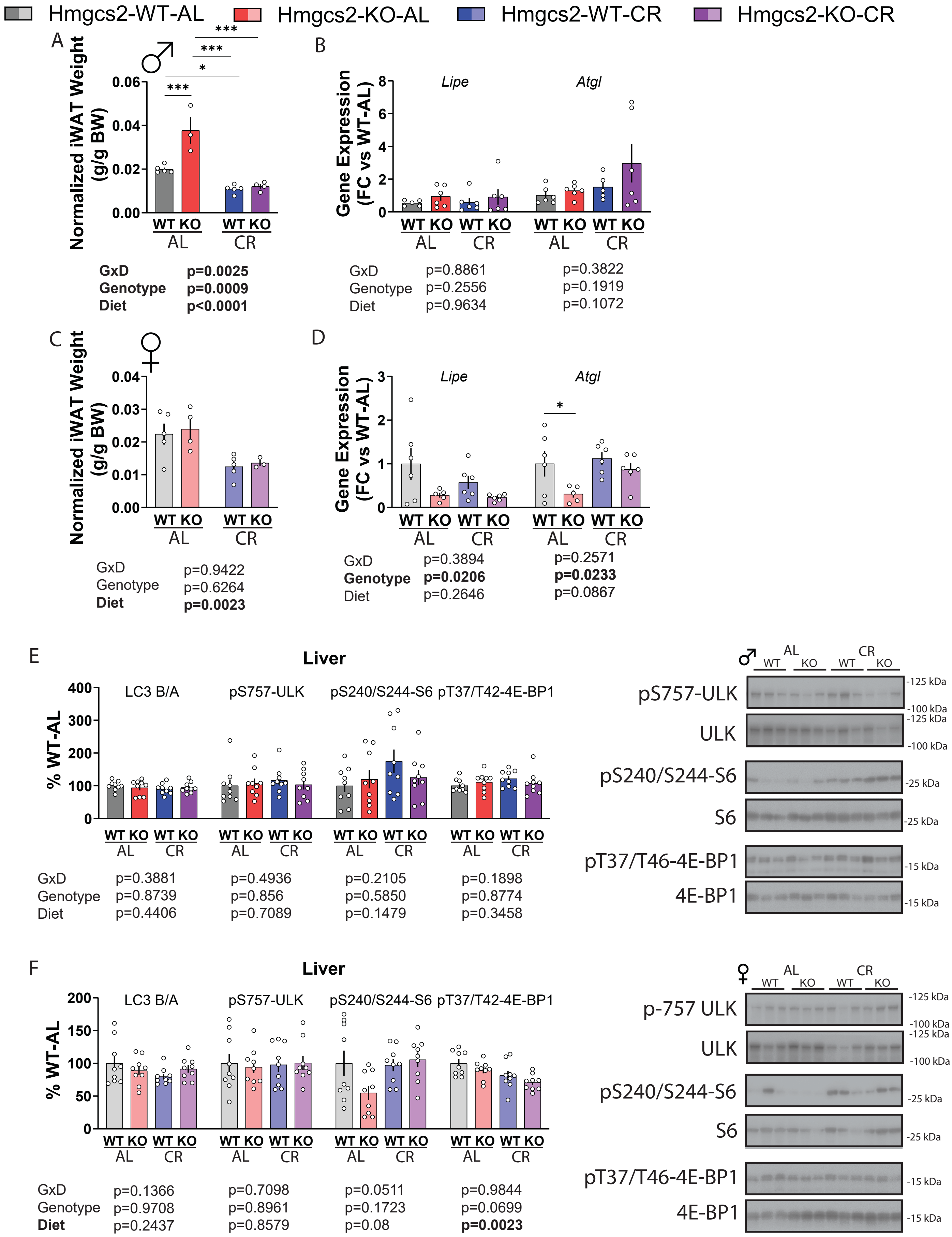

**Supplemental Figure 5. Lipolytic gene expression in the iWAT and mTORC1 signaling in the liver.**

**(A-B)** In male mice, body weight-normalized iWAT weight (A) and the expression of *Lipe* and *Atgl*. (B) in the iWAT of male mice. For WT-AL, KO-AL, WT-CR, and KO-CR, iWAT weight n=5,3,5,4; *Lipe* n=5,6,6,6; *Atgl* n=6,6,5,6. **(C-D)** In female mice, body weight-normalized iWAT weight (C) and the expression of *Lipe* and *Atgl* (D) in iWAT of female mice. For WT-AL, KO-AL, WT-CR, and KO-CR, iWAT weight n=5,4,5,3; both genes n=6,5,6,6. **(E-F)** LC3B/A ratio and mTORC1-related protein phosphorylation in male (E) and female (F) mice. n=9 each group. (A-F) *$p<0.05$; Sidak's test post 2-way ANOVA conducted separately for each gene (A-D) or protein/phosphor-residue (E-F). Data presented as mean ± SEM.

Supplemental Figure 6.

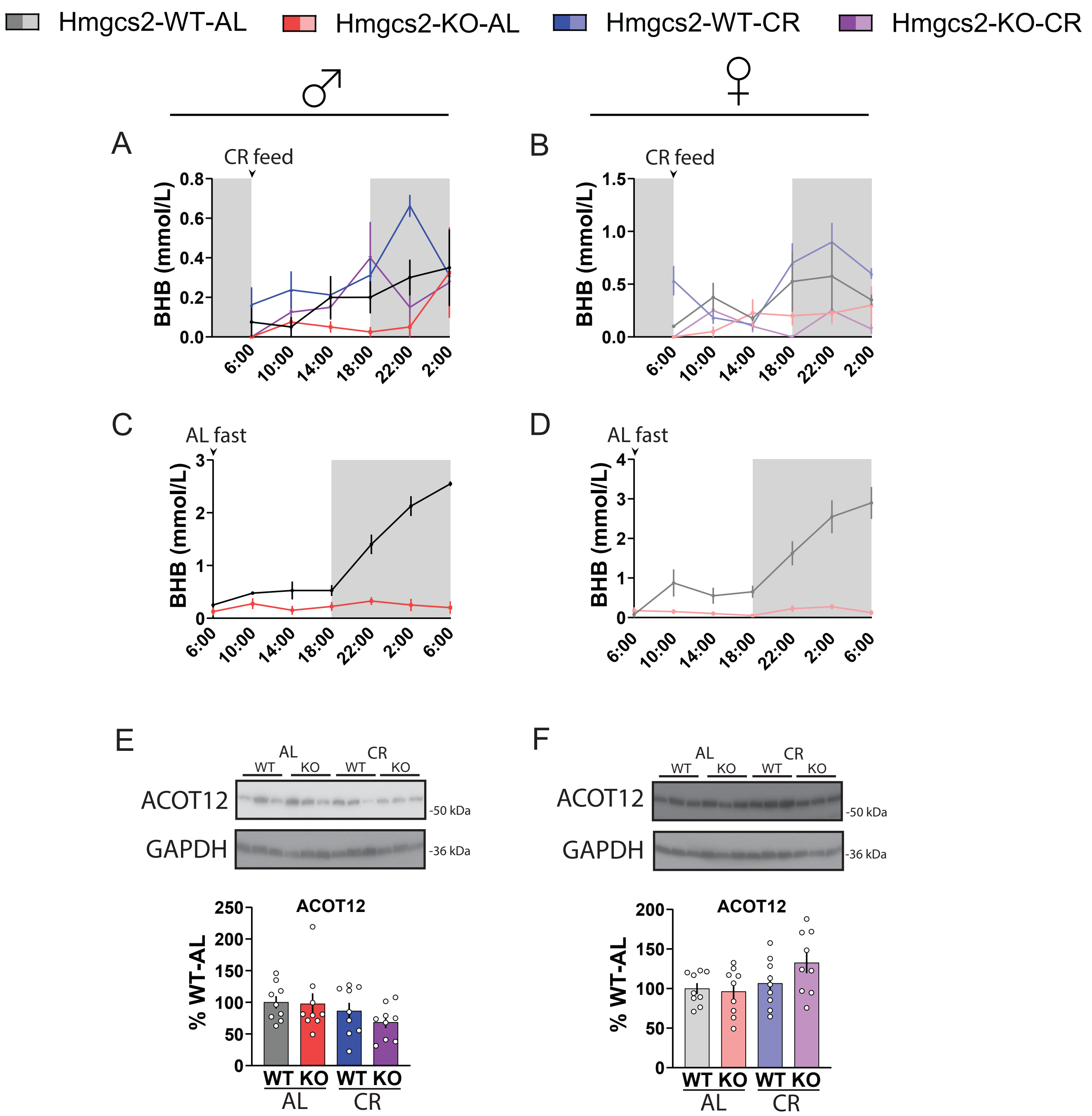


GxD p=0.5374
Genotype p=0.0419
Diet p=0.419

GxD p=0.1599
Genotype p=0.2908
**Diet p=0.0454**

**Supplemental Figure 6. Circulating βHB profile of animals during experiments and fasting; liver ACOT12 levels.**

**(A-B)** Circulating βHB level of male (A) and female (B) mice throughout a typical day. For WT-AL, KO-AL, WT-CR, and KO-CR, male n=4,4,8,4; female n=4,4,6,4. **(C-D)** Circulating βHB level of AL-fed male (C) and female (D) mice while fasting. n=4 each group. **(E-F)** Liver western blots in male (E) and female (F) mice for ACOT12. n=9 each group. Sidak's test post 2-way ANOVA. Data presented as mean ± SEM.
